# GlycoDiveR: a modular R framework to analyze and visualize highly dimensional glycoproteomics data

**DOI:** 10.64898/2026.03.21.713336

**Authors:** Tim S. Veth, Nicholas M. Riley

## Abstract

Mass spectrometry-based glycoproteomics is a critical platform for understanding the complex roles of protein glycosylation in biological systems, yet visualizing multidimensional glycoproteomics datasets remains a significant bottleneck in data interpretation and communication. Glycan microheterogeneity, i.e., the potential for a glycosite to be modified by multiple glycans, defies the binary presence-absence logic used in analyses of other post-translational modifications. Instead, glycoproteomics necessitates intentionally designed data structures and visualizations that are glycoform-centric, not just site-centric. Additionally, there is a need for complementary degrees of data analysis that alternate between glycoproteome-scale patterns and glycosite-specific regulation. Several bespoke frameworks for visualizing glycoproteomics data have emerged, but they often require advanced programming expertise and are designed for a single study rather than broad application. Here, we present our efforts to harmonize post-search data analysis of glycoproteomics through a modular R framework called GlycoDiveR. This platform streamlines import, transformation, and curation of qualitative and quantitative glycopeptide identifications, including support for raw output from multiple search engines. GlycoDiveR is designed to integrate seamlessly into existing analysis workflows by enabling fast, flexible exploration of highly dimensional glycoproteomics datasets via a consistently formatted data architecture. Our goal is to offer a customizable set of glycosylation-specific visualizations with minimal coding, while keeping data accessible to users who wish to further customize their characterization strategies. It also maintains a modular design that supports the continual addition of visualizations, analyses, and export functions. Ultimately, GlycoDiveR is meant to improve accessibility of glycoproteomic-specific analyses and lower the barrier to exploring biological narratives embedded in rich glycoproteomic datasets. GlycoDiveR is open-source and freely available at https://github.com/riley-research/GlycoDiveR.

## INTRODUCTION

Consistent advances in mass spectrometry (MS) instrumentation and related technologies^1–4^, sample preparation and enrichment strategies^5–7^, and informatics tools designed for glycopeptide identification^8–10^ have translated to an impressive expansion of scale and breadth in glycoproteomics. Indeed, it is becoming routine to report thousands to tens of thousands of glycopeptide identifications in a given experiment.^11–17^ Despite these advances, bridging the gap between search engine output and biological interpretation remains a challenge. Glycoproteomics as a field is still in a developmental phase, and many obstacles revolve around standardizing rigor in data analysis, including identification, quantitation, and how to interrogate the multiple dimensions of information encoded in glycopeptide data.^18–21^ The manifold heterogeneity of the glycoproteome is a unique feature that contributes to these challenges. Protein glycosylation is the covalent modification of proteins by mono- and polysaccharides (i.e., glycans), with substantial variation in glycan composition and structure that depends on the glycosylated amino acid, cell state, tissue of origin, and the organism overall.^22–24^ Each glycosylation site can be described in terms of its macroheterogeneity (i.e., the binary modified/non-modified state fundamental to all post-translational modifications [PTMs]) and microheterogeneity (i.e., the variety of glycans that modify any one site).^25^ The biological implications of each glycosite depend on both its macro- and microheterogeneity^26–28^, requiring the ability to sift through large-scale glycoproteome datasets with glycosite-level resolution.

Most glycoproteomics data analysis tools have built on progress and platforms used for other PTMs (e.g., phosphoproteomics^29–33^), but the binary presence/absence logic sufficient for interrogating those PTMs provides an inadequate framework for exploring multi-faceted glycoproteomics datasets. Instead, visualization approaches that embrace glycoform heterogeneity are critical to understanding the complexity of the glycoproteome. The mismatch between this need in glycoproteomics data analysis and the tools available has been well-recognized in the field.^34–38^ To circumvent a lack of glycoproteome-centric tools, numerous creative visualizations have been developed to interrogate glycoproteomics data at the glycosite and global glycosylation levels.^39–53^ These analyses are often custom-designed for a specific study, meaning efforts needed to generate them are not always straightforward to translate to new experiments or to recreate across different laboratories. Even so, each visualization typically builds from an approach that includes: a) compiling glycosylation-specific data from search engine outputs, b) cleaning and converting data in a manner that includes quantitative data and handles diverse glycan compositions while accommodating other non-glycan modifications, c) integrating glycosylation-specific annotations [e.g., assignment of glycan classes], d) formatting data to fit architectures needed for a visualization, and e) actual plotting of data and manipulation of display characteristics like colors, labels, and groupings. While these steps are readily achievable for tailored situations, the lack of an integrated platform to accomplish these tasks makes routine visualization of glycoproteomic data challenging and accessible only to users with advanced programming skills. Beyond limiting our collective ability to explore biological implications in glycoproteomics data, the lack of a streamlined workflow also hinders reproducible analyses and visualization strategies across the field. Importantly, data exploration is typically the most time-consuming step in glycoproteomics workflows, which means that a lack of workflow translates to slower project turnaround and longer times needed between experimental iterations.

To address this gap in post-identification visualization for the complexity of glycoproteomics data, we developed GlycoDiveR as an open-source, modular R framework that enables users to qualitatively and quantitatively assess glycoproteomie-scale and glycosite-scale data with >25 customizable visualizations that can be generated with minimal programming expertise. GlycoDiveR is designed to integrate into existing analysis pipelines by supporting raw search-engine output and providing direct access to formatted data, ensuring compatibility with subsequent analysis steps. GlycoDiveR supports all classes of protein glycosylation and curates identified glycopeptides while allowing non-glycosylation modifications. Its modular design enables continuous integration of novel visualizations, thereby providing a framework for community-driven evolution. We aim for GlycoDiveR to be a step toward bridging the gap between search-engine output and biological interpretation through visualization.

## EXPERIMENTAL AND METHODS

### Implementation and usage

GlycoDiveR is an open-source visualization and analysis framework implemented in R that directly imports results from FragPipe (N- and O-glycoproteomic workflows), Byonic, and pGlyco search engines, as well as MSstats and Perseus statistical platforms. At its core, GlycoDiveR uses two standardized file formats: one for search-engine output and the other for comparison data (e.g., MSstats comparison output). GlycoDiveR’s import function performs all necessary filtering and formatting, is modular to accommodate user needs, and requires a single function call to prepare the data for all visualizations (**Figure S1**). All input is converted to a standardized GlycoDiveR format, enabling straightforward comparisons, combining results from different search engines, and, most importantly, providing direct compatibility with >25 customizable publication-quality visualizations and processing functions, each accessible with a single line of code.

Search engine importing requires three files: the raw search engine output, an annotation file generated by GlycoDiveR’s GetAnnotation function that specifies the experimental design, and the FASTA file used for the glycopeptide search. This enables the transition from search engine output to the first visualizations to happen within minutes. Using the modular search engine import function, glycoproteomic data are filtered and transformed via data retention and normalization approaches that the user can select and evaluate. The resulting GlycoDiveR data is an R list object that includes PSM- and PTM-level data, annotated with features such as glycan categories and UniProt information^54^, including subcellular localization and domain annotations, and GyTouCan identifiers.^55–57^ UniProt information is retrieved through the UniProt website API.^58^ GlyTouCan identifiers are retrieved by converting glycan compositions to WURCS,^59,60^ which are subsequently converted to GlyTouCan identifiers; both conversions are performed using the GlyCosmos API.^61,62^ GlycoDiveR data also includes the combined raw search engine output with an identifier linking all PSMs to the other data frames, allowing users to verify the filtering and data organization and to add custom columns. Data within the GlycoDiveR format (i.e., in an R list object) are stored as data frames, allowing users to easily access and modify them as in other R workflows. Users can customize sample grouping and definitions, colors, and other data details as needed. A valuable feature of GlycoDiveR is the ability to explore subsets of data (e.g., [glyco]peptide, protein, and sample levels) without modifying the main data frame. These subsets can be used in all visualizations via the whichPeptide, whichProtein, and whichAlias arguments. By default, unnormalized and normalized quantitative values are imported, and can be updated using output from MSstats,^63^ Perseus,^64^ or custom tables. GlycoDiveR can also perform comparisons using two-tailed t-tests and the Benjamini-Hochberg correction to obtain adjusted p-values.^65^ Although this is a relatively straightforward and statistically sound approach, GlycoDiveR is not intended to replace specialized statistical packages such as Limma or MSstats,^63,66^ but rather to be integrated into workflows that leverage these statistical packages prior to data visualization.

Glycopeptides are automatically categorized by glycan type during data import. The default categorization for N-glycans is determined in the following order, so that the first categorization that fits the glycan is used and no subsequent categories are assigned: contains sialic acid and fucose = Sialofucosylated; contains sialic acid = Sialylated; contains fucose = Fucosylated; contains glycan phosphorylation = Phosphomannose; any structure smaller than or equal to the N-core (N(2)H(3)) = Truncated; contains fewer than 3 HexNAc residues and more than 3 hexose = Oligomannose; otherwise, it is categorized as a Complex/Hybrid glycan. Glycans attached to serine or threonine are categorized as O-glycans. Our group largely focuses on mucin-type O-glycosylation over O-GlcNAc, but users can create specific O-glycosylation categories as needed. All other glycans are categorized as non-canonical glycans. GlycoDiveR does not use a paucimannose N-glycan category, as these are typically reserved for glycans generated by hexosaminidases, which cannot be deduced solely from glycan composition.^67^ Therefore, ‘Truncated’ is used for these glycan compositions. Users can create their own categorization definitions and apply them within the GlycoDiveR framework to override default categories as needed.

### Datasets used in this study

To illustrate GlycoDiveR’s functionality, we re-analyzed data from Kawahara et al.^68^, which is publicly available through the ProteomeXchange Consortium via the PRIDE partner repository with the dataset identifier PXD051882.^68,69^ The data consist of four TMT-labeled glycopeptide pools, each containing samples from one stage of colorectal cancer, the corresponding adjacent tissue, and a pooled TMT reference. Data was re-analyzed using FragPipe (v24.0)^70^, using the Glyco-N-TMT workflow, and the reviewed human protein database download from Uniprot (January 2026)^54^ to which decoys and contaminants were added as part of the FragPipe workflow. Default settings were used unless specified otherwise. In short, the search was performed using a 20 ppm precursor mass error and fragment mass tolerance, two allowed miscleavages (KR), a maximum of 3 variable modifications (M oxidation, N-terminal acetylation), and fixed modifications set to carbamidomethylated C and TMT-labeled N-terminus and lysine. The ‘Human_N-glycans-medium-253` glycan database was used with a 1% glycan FDR. TMT10-plex was selected using MS2-level quantification. PSMs output files are available as supplemental material. Samples were labeled as specified in Kawahara et al. All (glyco)PSMs were filtered at 1% FDR (q-value ≤ 0.01 and glycan q-value ≤ 0.01) using GlycoDiveR.

## RESULTS AND DISCUSSION

We distilled years of developing glycoproteomics-centric data analysis and visualization into an open-source R framework called GlycoDiveR. This platform allows users to rapidly assess data quality and to explore biological narratives in glycoproteomic data through glycosylation-specific visualizations. The philosophy of GlycoDiveR is multi-fold: 1) to provide a framework for examining technical quality and underlying biological implications in glycoproteomics data through a highly customizable set of glycosylation-specific visualizations; 2) to standardize a platform for comparing glycoproteomics workflows used by a variety of research groups; 3) to make these analyses accessible to both glycoproteomics experts and non-experts with minimal coding required while also maintaining a flexible architecture for domain experts who want to further customize visualizations and analyses; and 4) to maintain a modular design that supports the continual addition and refinement of visualizations, analyses, and export functions. Streamlining these processes shifts the focus of glycoproteomics data analysis from a computational burden associated with data wrangling and poorly defined data architectures to an ability to rapidly evaluate results and move toward biological interpretation. Our goal is that GlycoDiveR will be used by glycoproteomics-focused labs, but also as a platform to share data with non-domain-expert collaborators to empower them to explore biological insights embedded in information-rich glycoproteomics datasets.

At the heart of GlycoDiveR lies the standardized GlycoDiveR format that allows direct compatibility with all GlycoDiveR functions. All raw search engine output is converted into this format using a single GlycoDiveR function, for example, ImportMSFragger(), which provides full control over any filtering, annotation, and formatting of the data (**Figure 1**). To showcase how GlycoDiveR powers the analysis of glycoproteomic data, we re-analyzed data from Kawahara et al. (PXD051882), which comprises four TMT pools, each including samples from stages 1-4 colorectal cancer (CRC), adjacent healthy tissue, and a pooled TMT channel. The pooled TMT samples were used to normalize intensities between the four pools. Note, all presented visualizations in this manuscript (other than Figure 1) were generated by GlycoDiveR with minimal organization in Inkscape.^71^ We intentionally kept scaling, colors, and labeling as they appear from GlycoDiveR outputs to underscore the publication-quality graphics GlycoDiveR exports. Not all available visualizations can be shown here, but all visualization functions have arguments to modify the aesthetics and non-aesthetics e.g., how the underlying data are represented or summarized) of the visualization. All functions are described in detail on GlycoDiveR’s GitHub: https://github.com/riley-research/GlycoDiveR.

**Figure 1.**
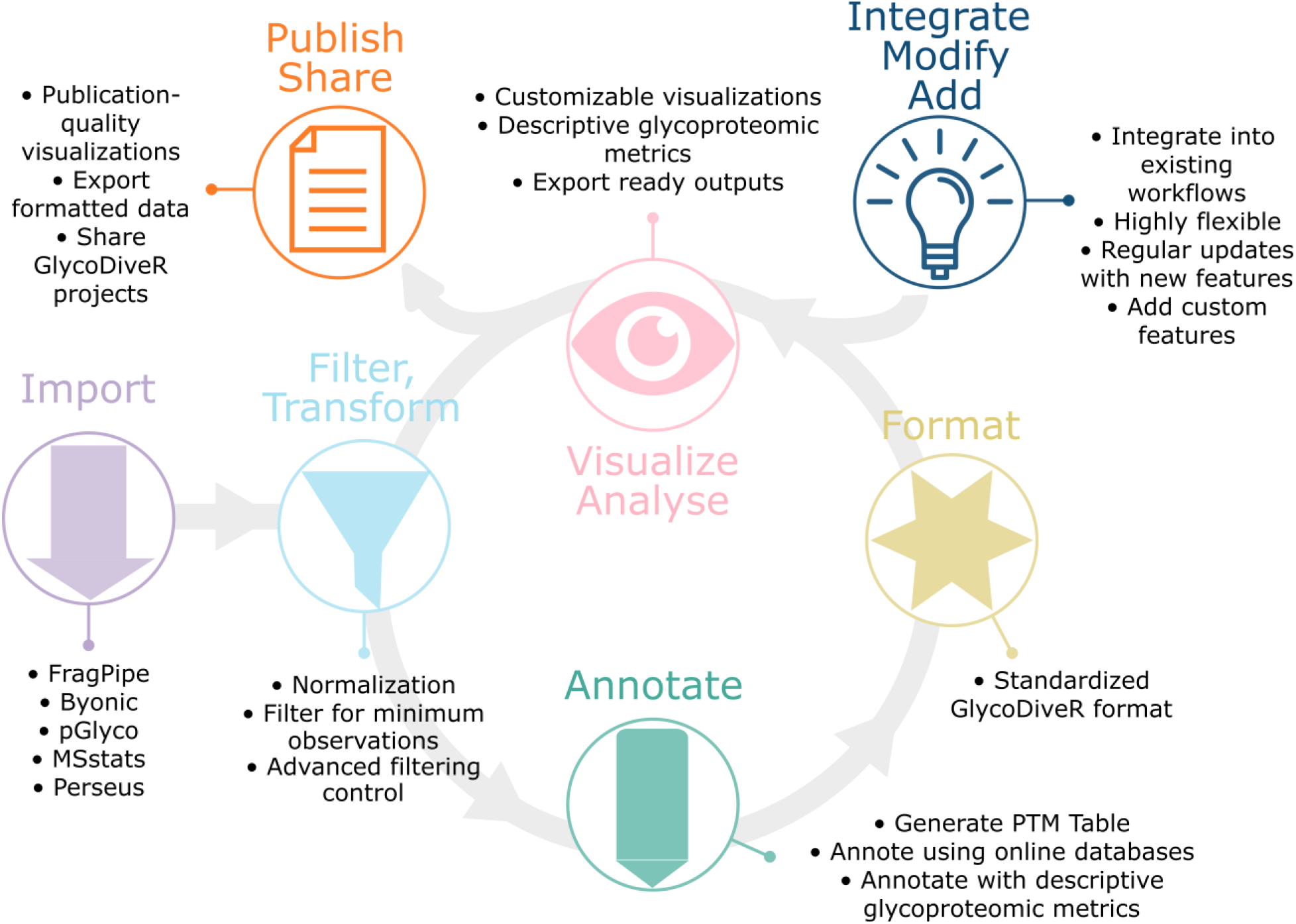
The GlycoDiveR workflow. GlycoDiveR imports, filters, transforms, annotates, and formats raw search engine output into a standardized GlycoDiver format. All of these steps are completed in a single function call. GlycoDiveR architecture ensures that users have full control over the data and that the imported data is compatible with all downstream functions. The circular workflow here illustrates how GlycoDiver enables iterations of data analysis, visualization, and re-analysis based on insights learned through visualization. Data frames and publication-quality visualizations can be exported directly from GlycoDiveR, and the modular design ensures that new visualizations can be easily incorporated into the workflow in future iterations.

### Verifying sample and experimental quality with GlycoDiveR

Search engine performance remains a challenge in glycoproteomics^18^, so most workflows use additional filtering steps to remove low-quality data. A unified strategy or general set of metrics has yet to be identified by the field, and useful filtering metrics often differ by search engine. GlycoDiveR provides flexible user-defined decisions for filtering search engine results to keep the platform broadly compatible with multiple workflows. For the MSFragger-Glyco search used in our re-analysis here, we filtered for a 1% FDR at the peptide and glycan levels (q-values ≤ 0.01). For certain visualizations as indicated below, we filtered for glycopeptides with a CV < 20% or with at least 70% of quantitative values in at least one group. GlycoDiveR allows filtering for any criteria in the search engine output, so users can choose metrics suitable for their experiments, and data analysis with GlycoDiveR can adapt to best practices as they are refined by the field.

Comparing identifications for qualitative evaluations is a standard first step in most workflows, which GlycoDiveR enables at the PSM and unique (glyco)peptide level (**Figure S2**). Common qualitative evaluations, like peptide length distributions and mass error versus retention plots, are also available (**Figure S3**). Additionally, technical quality is central to the reliable extraction of biological insights, with evaluations needed to ensure accurate sample preparation, control of batch effects, and proper data normalization,.^72,73^ Each dataset is unique; therefore, normalization and filtering methods must be evaluated for each experiment. Inappropriate normalization and filtering schemes may prevent the extraction of differentially regulated glycoforms or introduce false positives among the glycoproteins identified as differentially expressed.^74^ GlycoDiveR always retains both the raw and the normalized intensity values to facilitate these validations and includes comparison plots (e.g., box plots of intensity distributions) to validate normalization strategies (**Figure 2a**). Users have several options for normalization they can evaluate (**Figure 2b**), providing flexibility to select the method most appropriate for each experiment. PCA plots of the Kawahara data (**Figure 2c** and **2d**) illustrate the importance of verifying normalization strategies. Here, median normalization led to clustering by TMT pool, a classic indication of batch effects that could lead to false positives or false negatives when evaluating differentially expressed glycopeptides.^75^ Thus, in this example, raw intensity is a better strategy than median normalization for further quantitative processing. **Figure S5** shows an example of label-free quantitative data from IPX0011732000^76^ where median normalization does not induce batch effects and is a reasonable choice for further quantitative analysis. GlycoDiveR makes this decision straightforward to evaluate, helping to avoid potentially critical mistakes that affect downstream conclusions. GlycoDiveR also generates loadings plots for further evaluation of PCA clustering (**Figure S4**). As an additional quality control, GlycoDiveR also generates sample-specific or whole-dataset distributions of glycopeptide coefficients of variation (CVs) using either regular CV calculations on non-transformed intensity data or geometric CV calculations on transformed intensity data^77^, enabling filtering to retain glycopeptides that meet user-specified CV criteria (**Figure S6**).

**Figure 2.**
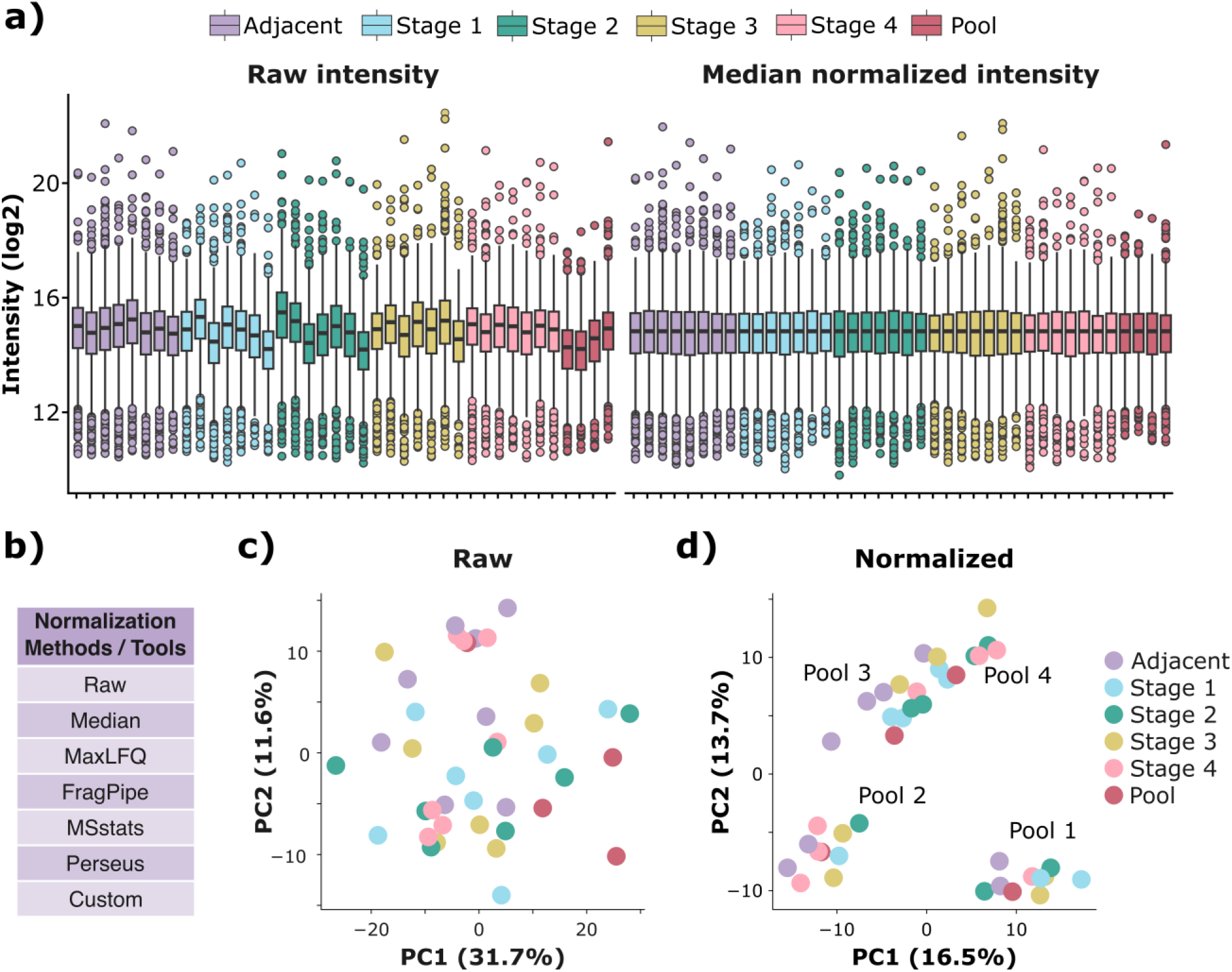
Verifying the technical quality of glycoproteomic data. (**a**) GlycoDiveR can perform several data normalization tasks and stores raw and normalized quantitative values. The PlotAbundanceQC function allows visualization of both intensity values and the effects of normalization. The left panel shows raw intensities from the TMT quantitation provided by data processing steps prior to GlycoDiveR import and the right panel shows median normalized intensities. (**b**) A list of supported normalization methods and tools that can be selected with a single argument to the importer function. PCA plots of samples with (**c**) raw intensity values and (**d**) median-normalized data reveal batch effects that occur with median normalization, a common side effect of isobaric labeling strategies. Each ‘Pool’ label in the right PCA plot with normalized data was added outside of the GlycoDiveR environment to show where a single pooled TMT sample is clustered; these labels are not included in the PCA using raw intensities because pools were not clustered. Note, Figure S5 shows a more typical example of label free data where median normalization does not create batch effects.

### Glycoproteome-scale Visualizations

After initial quality control, exploring glycoproteomics data can be done at multiple levels (e.g., whole dataset, subsets of glycoproteins, specific glycosites). Often it can be advantageous to visualize “glycoproteome-scale” trends that incorporate the whole dataset to begin interrogating broad biological narratives that emerge from the experiment. This requires numerous graphs that incorporate various layers of annotation, including glycan categories derived from composition information, multiple degrees of overlap between data sets (e.g., glycans, glycosites, glycoproteins), and glycan-protein connectivity that must be accounted for at each level of analysis. A single visualization rarely tells the full story, so we designed GlycoDiveR to offer a diverse portfolio of visualizations that can be quickly graphed, assessed, and customized.

A common comparison in glycoproteomic studies is differential expression analysis of glycopeptides. GlycoDiveR imports comparison results from MSstats,^63^ Perseus,^64^ or custom analyses and converts them to GlycoDiveR’s comparison table format. The comparison table, which can also be computed internally within GlycoDiveR, contains log2 fold changes and statistical metrics, including p-values and multiple-comparison-adjusted p-values from one or multiple group comparisons. The internally computed comparison table uses two-tailed t-tests and a Benjamini-Hochberg FDR correction; however, if more advanced statistical control is needed for an experiment, we recommend using tools such as MSStats and Limma prior to GlycoDiveR import.^63,66^ The comparison table can be used to generate volcano plots, which are customizable through user-defined selection of p-values or adjusted p-values, significance thresholds, glycopeptide labeling settings, and modifiable color schemes. **Figure 3a** displays a volcano plot with comparisons internally computed by GlycoDiveR for stage 3 CRC versus adjacent tissue. Labels show up- and down-regulated glycopeptides from multiple proteins known to be involved in CRC, such as integrin subunit alpha 6 (ITGA6)^78^ and carcinoembryonic antigen-related cell adhesion molecules (CEACAM)^79^. Glycopeptide subsets can be readily extracted and used in any GlycoDiveR visualization, such as the bar graphs in **Figure 3b** that show normalized intensity contributions by various glycan categories for up- and downregulated glycopeptides. Kawahara et al. reported an increase in truncated glycans in later stages of CRC, which is consistent with data generated here by GlycoDiveR. Glycoproteins can also be ranked by abundance in Glycoprotein Rank plots to understand how their relative abundances in samples shift relative to other conditions (**Figure S7**).

**Figure 3.**
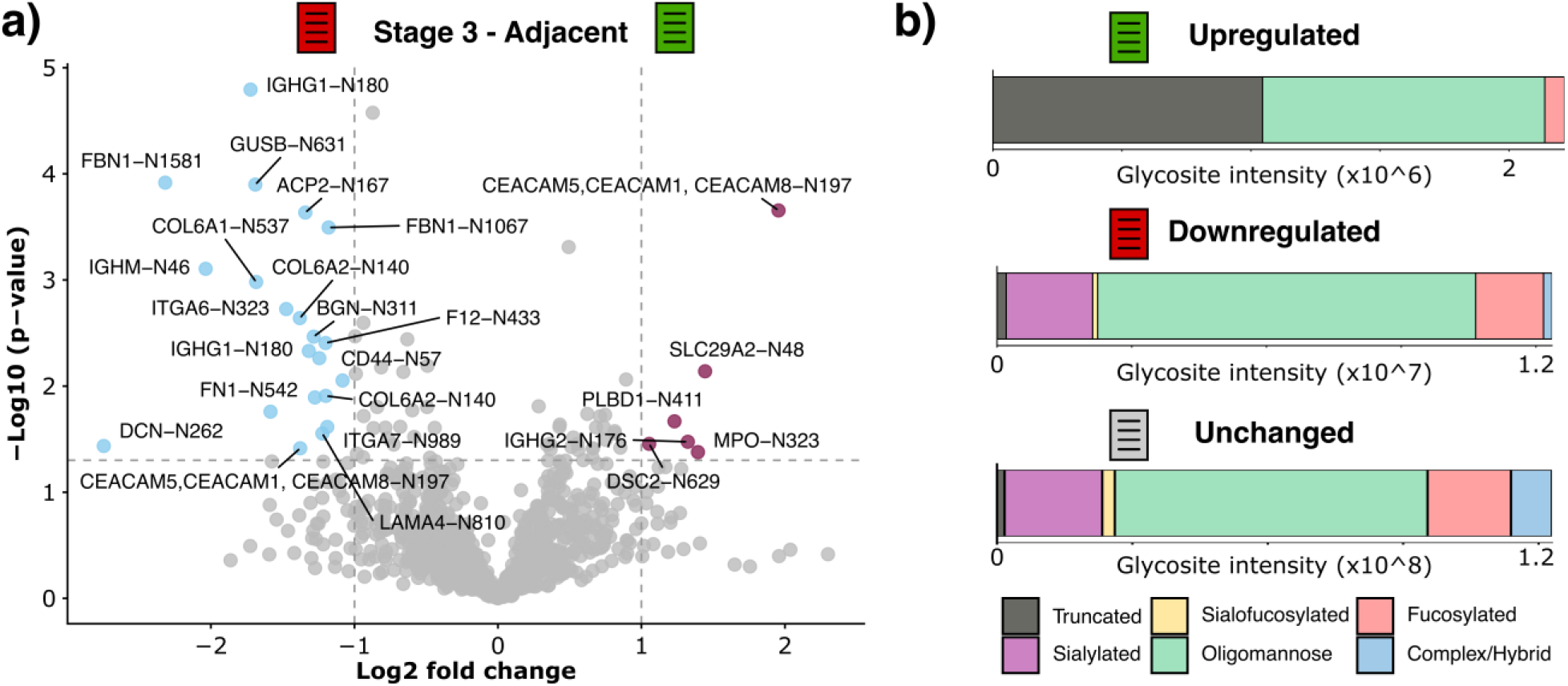
Visualizing differential regulation of glycopeptides. (**a**). A volcano plot showing fold changes for glycopeptides quantified in stage 3 CRC relative to adjacent tissue. Each point is a glycopeptide, with labels provided by GlycoDiveR for significantly regulated species. Log2 fold changes and p-values were internally computed in GlycoDiveR. (**b**) The significantly up- (green icon, right side of volcano plot) and down-(red icon, left side of volcano plot) regulated glycopeptides were extracted with a filtering function in GlycoDiveR, and their glycan compositions were visualized based on contributions to total glycopeptide intensity. Glycopeptides with truncated glycans comprise ∼50% of the signal of upregulated glycopeptides (top) but only ∼ 2% of the signal of downregulated glycopeptides (middle). Glycan categories can also be graphed for unchanging glycopeptides for comparison (bottom).

One useful, but less standard visualization we have developed through recent work^80^ is a Completeness Matrix that shows quantified and non-quantified/missing glycopeptides across conditions, providing an overview of commonalities across samples. GlycoDiveR generates Completeness Matrices of unique glycopeptides grouped by glycan categories and enables filtering to retain species identified in user-defined percentages of conditions (**Figure 4a**). Users can easily visualize how adjusting the minimum requirements to retain glycopeptides affects dataset completeness. This can be a vital step for further data curation, as glycoproteomic experiments can suffer from high levels of missing values that must be carefully managed.^81–83^ Note, imputation occurs prior to import into GlycoDiveR (e.g., with MSstats), so Completeness Matrix filtering should reflect decisions made in prior steps. GlycoDiveR also generates Venn Diagrams and UpSet plots to compare shared and unique identifications (**Figure S8**).

**Figure 4.**
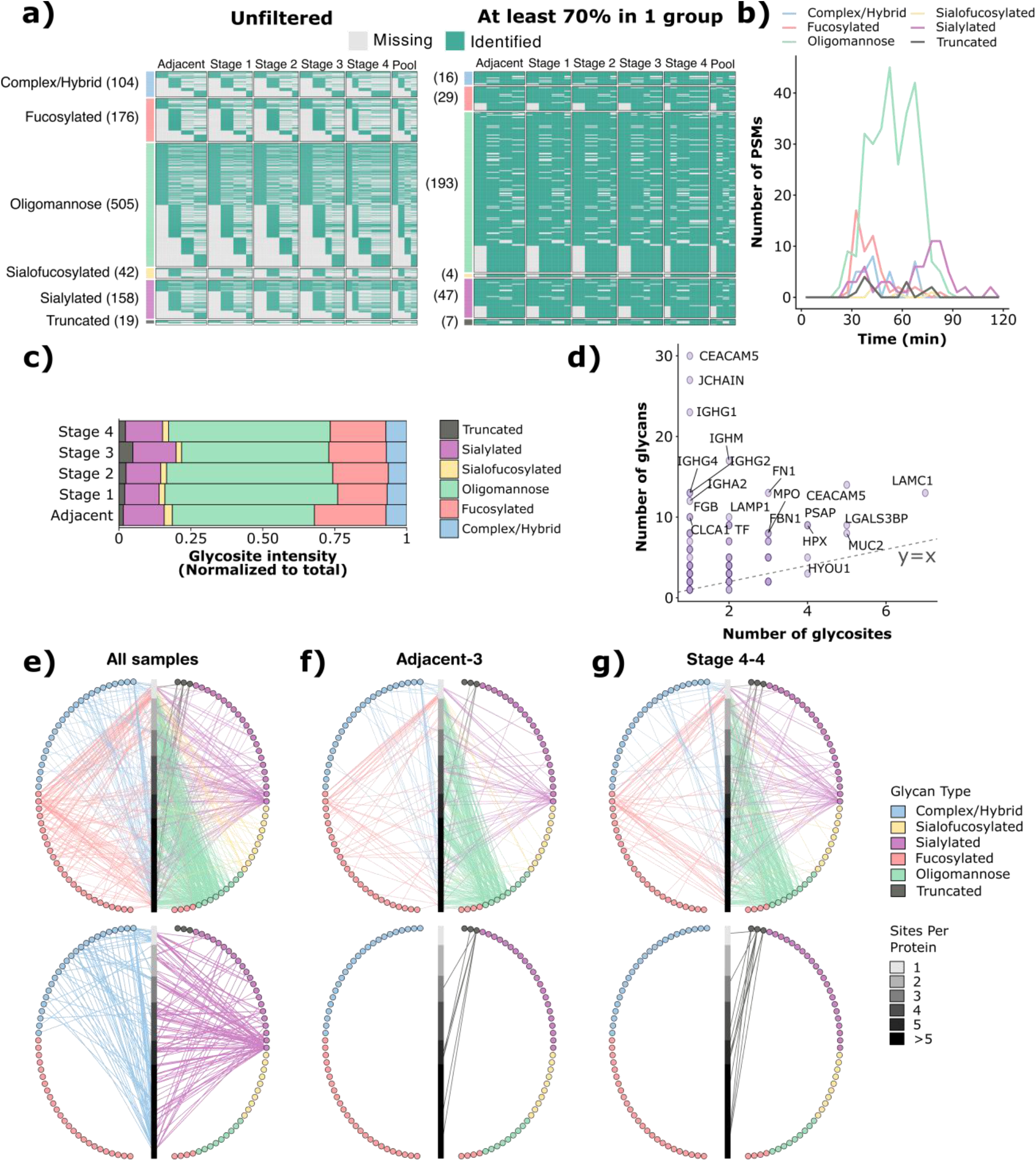
Glycoproteome-scale plots capture trends across complex glycoproteomics data. (**a**) Completeness Matrices show quantified (green) or non-quantified/missing (grey) glycopeptides to visualize dataset completeness. Glycopeptides are sorted by glycan category, as shown by colored bars to the left of each group in the matrix. Numbers next to each glycan category show the total number of species in each group. The two different matrices show the Kawahara 2025 dataset imported without filtering for a minimum number of observations (left, n = 1004 N-glycopeptides) and with filtering that required quantitative values for at least 70% replicates at least one group (right, n = 296 N-glycopeptides). (**b**) GlycoPSMs are grouped by their glycan category and graphed as a function of retention time. A bin width of 5 minutes is used in this example, but this value can be changed with a single argument in the plotting function. (**c**) Intensities of all quantified glycopeptides were summed per glycan category and normalized to show relative proportions across each sample (raw summed intensities can also be graphed with this function in GlycoDiveR). This analysis revealed an upregulation of truncated glycans, particularly in stage 3 CRC, corroborating the results of Kawahara et al. (**d**) The Glycans-versus-Glycosites Scatter Plot shows the number of unique glycan compositions detected for a protein (y-axis) graphed as a function of the number of identified glycosites for that protein (x-axis), revealing glycoproteins with particularly high or low microheterogeneity. (**e**) GlycoDiveR generates Glycoprotein-Glycan (GPG) networks that serve as a fingerprint for the entire glycoproteome detected in the experiment (top, all samples in aggregate). Specific glycan features can also be mapped across the full experiment, e.g., sialylated and complex/hybrid glycans detected in all samples (bottom). Panels (**f**) and (**g**) show GPG networks for Adjacent-3 and CRC Stage 4-4 samples, respectively. The top GPG networks show all glycopeptides detected in each sample while the bottom networks show only the truncated glycans.

Grouping glycopeptides by glycan categories can also be useful for evaluating how experimental parameters effect outcomes, such as visualizing if the LC gradient appropriately distributed glycopeptide identifications across retention times (**Figure 4b**). Insights from these graphs can be used for iterative experiment design to optimize conditions for specific glycopeptides of interest. Fucosylated glycopeptides, for example, eluted primarily in the first half of the chromatogram, indicating that a shallower gradient slope could potentially improve fucosylated glycopeptide characterization. **Figure 4c** shows how visualizing the abundance of glycan categories across the entire glycoproteome can reveal global trends, such as an increase in the abundance of truncated glycans in later stages of CRC. This again aligns with the observations of Kawahara et al., who demonstrate upregulation of truncated glycans in CRC due to increased expression of N-acetyl-β-D-hexosaminidase (Hex) subunit β (HEXB).

Other glycoproteome-scale visualizations attempt to summarize the degree of heterogeneity with some insights into protein-glycan combinations. One such example developed in Riley et al.^41^ is a Glycans-versus-Glycosites Scatter Plot, in which each point designates a glycoprotein (**Figure 4d**). The number of total glycosites observed for a protein is along the x-axis, while the number of total glycan compositions seen to modify glycosites of the that protein is along the y-axis. If a protein had exactly one unique glycan at each glycosite, it would fall on the y = x line. Glycoproteins with a high or low degree of microheterogeneity are above or below this y = x line, respectively. Another unique glycoproteomics visualization developed by Riley et al.^41^ is the Glycoprotein-Glycan (GPG) network. This bipartite network strategy has been adopted by several groups^51,81,84–87^, so we enabled GPG networks as a visualization within GlycoDiveR (**Figure 4e-g**). GPG networks function as a “fingerprint” of the glycoproteome when viewing an entire dataset, but they can also be filtered to show how specific glycan features are distributed across the glycoproteome (**Figure 4e**). They can be further used for comparing glycoproteome features between conditions, as shown in **Figures 4f** and **4g** that compare whole glycoproteome networks and truncated glycan features between adjacent tissue and stage 4 CRC samples. The GPG networks complement observations from Figures 3b and 4c that show upregulation of glycopeptides with truncated glycans in CRC. The GPG networks show that increases in truncated glycoforms in CRC come from a few glycan compositions primarily found on multiply glycosylated proteins without an increase in the number of truncated glycan compositions themselves.

Beyond visualizations discussed above that are largely self-contained by data in the search engine results, external databases and literature mining contribute additional value to data interpretation. GlycoDiveR currently interfaces with the Uniprot API^54,58^ to extract subcellular localization and protein domain annotations as part of the data import process, which it then incorporates into the GlycoDiveR standardized format for simple access and customization. GlycoDiveR also uses the GlyCosmos API^61^ to fetch GlyTouCan accession numbers to harmonize glycopeptide identifications with curated glycan-specific databases.^55,61,88^ The flexibility of GlycoDiveR’s design means additional information from either API connection, or additional databases in general, could be incorporated in future versions. **Figure 5** displays the number of glycopeptides detected per subcellular localization to demonstrate how these annotations can be incorporated into glycoproteomic data analysis. Changes in glycosylation were identified across subcellular compartments (as annotated in Uniprot), including the plasma membrane, multiple organelle membranes, and secretory vesicles. Glycosylated sites with specific annotations can be readily extracted and used in other GlycoDiveR visualizations for further characterization.

**Figure 5.**
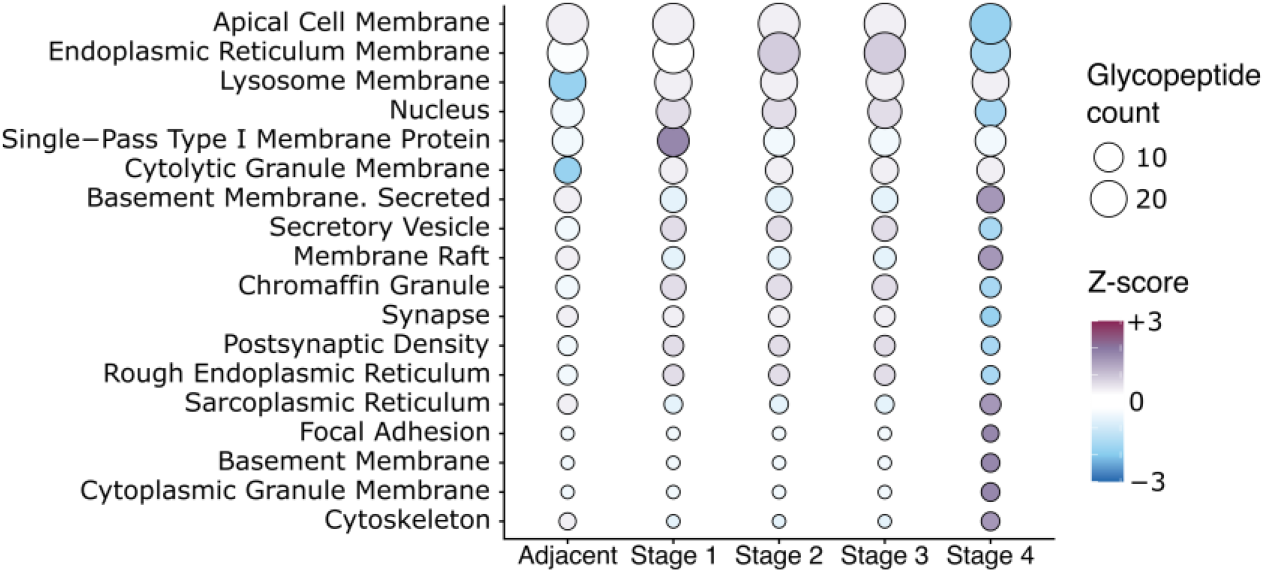
GlycoDiveR includes visualizations with annotations from online databases. The number of quantified glycopeptides is shown for each subcellular compartment, which is extracted from GO terms associated with their corresponding proteins in the UniProt database. Circles are scaled to the number of glycopeptides found and shaded based on the row z-scored values. Only compartments with a minimum Z-score of 1.5 are shown.

### Glycosite-scale Visualizations

Glycoproteome-scale visualizations described above are valuable for capturing global trends and revealing interesting subsets of data to investigate further. Narrowing the scope of interrogation to specific glycoproteins and glycosites provides a complementary approach to understanding a dataset. A key feature of GlycoDiveR is the ability to visualize glycoproteome- and glycosite-scale data in the same environment, and any visualization can be reduced from glycoproteome scale to a subset of data by providing filters for glycoproteins, glycopeptides, or samples of interest. A common example of where this “zoom in” on glycoproteins of interest can add value is understanding site-specific microheterogeneity across sample conditions. GlycoDiveR integrates domain information from Uniprot, quantitative information from FragPipe, protein length extracted from the FASTA file, and glycan classes to generate glycosite-centric plots for any glycoprotein in the dataset. For example, immunoglobulin M (IgM) is an interesting immune-response-relevant glycoprotein in the Kawahara dataset that has high glycosite microheterogeneity (i.e., it appears above the y = x in the Glycans-versus-Glycosites Scatter Plot in Figure 4d). **Figures 6a** and **6b** show Glycosite Maps of IgM glycosites from single replicates of adjacent and stage 4 CRC samples. These maps highlight the increased number of glycans (i.e., increased glycosite microheterogeneity) in the stage 4 CRC sample compared with adjacent tissue.

**Figure 6.**
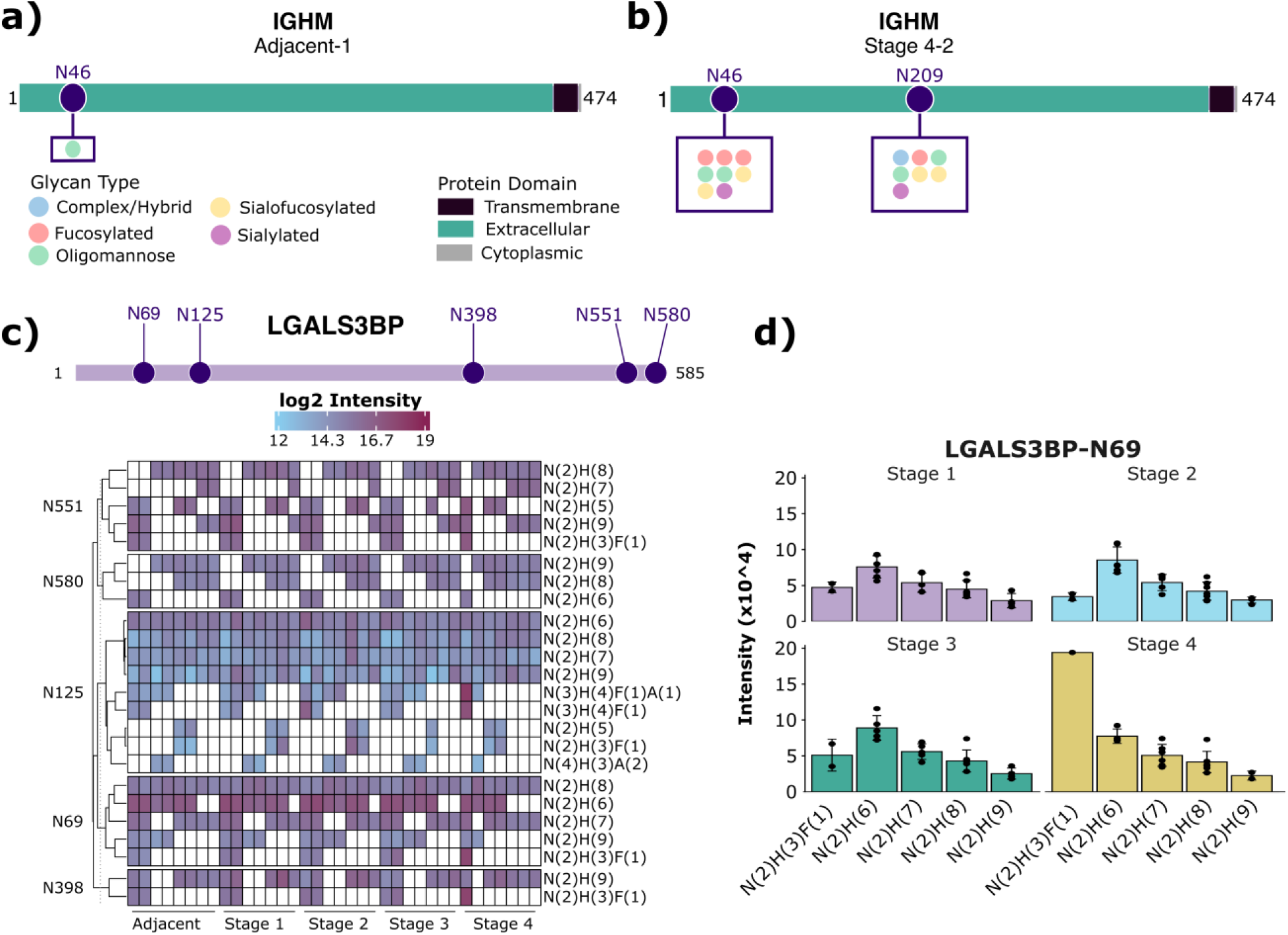
Glycosite-level visualizations. Integrating domain information from Uniprot, quantitative information from FragPipe, protein length extracted from the FASTA file, and glycan classifications in a glycoprotein-specific manner show distinct glycosylation profiles of IgM in (**a**) adjacent tissue replicate 1 compared to (**b**) stage 4 CRC replicate 2. (**c**) The quantitative heat map shows log2 glycosite intensities for the five detected N-glycosylation sites of LGALS3BP. White squares are non-quantified values. Rows were clustered per site using hierarchical clustering. (**d**) Quantitative data for glycoforms at a specific glycosite can be summed for each condition, as is shown here for LGALS3BP-N69 shown across the different stages of CRC. Error bars represent standard deviations. These graphs can be generated for any glycoprotein in the dataset.

Looking at quantitative values at the glycosite-level for a specific protein provides an additional layer of information, too. Galectin-3-binding protein (LGALS3BP) provides a demonstrative example; this glycoprotein is of putative CRC cell origin, is known to be involved in the glycoimmune axis for CRC,^89^ and was also a glycoprotein of interest for Kawahara et al. LGALS3BP showed a relatively large number of quantified glycosites and moderate-to-low glycoform heterogeneity (9 glycan compositions observed across the 6 sites),^89^ so we were curious how glycoform abundance of LGALS3BP varied at the glycosite level. The PlotPTMQuantification function in GlycoDiveR generates a quantitative heatmap for specific glycoproteins (**Figure 6c**), showing that LGALS3BP exhibits a sevenfold range in glycan abundance and consistent glycosylation across sample types. This heatmap makes it easy to see that oligomannose glycans are the most abundant and consistently identified glycan compositions for all glycosites, but that N125 sometimes harbors truncated and complex type glycans and N69 shows a highly abundant truncated glycoform in one stage 4 replicate. To get more granular for a specific glycosite, GlycoDiveR also enables quantitative barplots for individual glycosites for any glycoprotein in the dataset. Providing *whichProtein = “P01591”, site = “N69”*, and *whichAlias = “CRC”* arguments in the GlycoDiveR function PlotSiteQuantification generates the quantitative barplot in **Figure 6d** that reveals differences in glycoform levels across CRC stages. GlycoDiveR makes selecting samples and protein subsets for these types of graphs straightforward, so glycosite-level visualizations can be automatically generated and saved for any glycoprotein in the dataset with minimal coding required.

## CONCLUSIONS

GlycoDiveR aims to bridge the gap between search engine output and biological interpretation through data visualization specifically geared to the highly dimensional data encountered in glycoproteomics. Numerous useful visualization strategies have been created in prior studies to manage multifaceted glycoproteomics data, yet access to platforms to repurpose these graphs for other experiments have remained a limiting factor in analysis workflows. Instead, most of these prior efforts rely on bespoke, non-standardized data wrangling and advanced coding skills that create a disconnect in data analysis across the field. GlycoDiveR fills this gap by providing a diverse portfolio of visualizations accessible with minimal lines of code required. Importantly, the >25 visualizations that currently exist in the GlycoDiveR environment span both glycoproteome- and glycosite-scale visualizations, generating complementary strategies to decipher glycoproteomics data. We designed GlycoDiveR to integrate into current workflows that may vary across users, and its modular architecture can accommodate new visualizations and analyses as analytical strategies for glycoproteomics continue to mature. We aim to update GlycoDiveR frequently with new visualizations we develop through our glycoproteomics research (for example, incorporating other input data beyond glycopeptide search engine results^90^), but its open-source nature also means others in the glycoproteomics community can build new visualizations they develop into the GlycoDiveR environment. Additionally, GlycoDiveR extends beyond visualization by providing tools for data integration, exploration, and export. All data are formatted into hierarchical data frames to allow users to modify and explore the data with straightforward functions, and users can customize data and GlycoDiveR functions to meet their experimental needs. The platform also provides an avenue to extract descriptive statistics from glycoproteomic datasets, e.g., calculating the percentage of missing values in (a subset of) the data or computing the number of possible glycoforms of a protein. To extend beyond the GlycoDiveR platform, our data architecture enables standardized data export while also allowing users to write custom exporters. Data export from GlycoDiveR’s centralized data frame ensures consistent formatting and data annotation independent of the search engine used. This strategy not only enables exploration of the data by those who generate it, but it also facilitates sharing data with third party collaborators. Altogether, GlycoDiveR represents a step toward accessibility and reproducibility in glycoproteomic data analysis and lowers the barrier to exploring multiple layers of biological insights generated by modern glycoproteomic methods.

## Supporting information

TMT1_psm

TMT2_psm

TMT3_psm

TMT4_psm

## DATA AND CODE AVAILABILITY

GlycoDiveR is open-source and freely available on GitHub: https://github.com/riley-research/GlycoDiveR. The GlycoDiveR GitHub repository contains comprehensive documentation, an overview of the functionality, and user guides.

## SUPPORTING INFORMATION

The Supporting Information is available free of charge online.

- Figures in this manuscript used data from Kawahara et al. (PXD051882) that were reprocessed with MSFragger-Glyco. Search results that were imported into GlycoDiveR are available as supplemental files: TMT1_psm.tsv, TMT2_psm.tsv, TMT3_psm.tsv, and TMT4_psm.tsv
- Supplemental Figures include: Figure S1: Architectural overview of GlycoDiveR; Figure S2: Comparing identifications with GlycoDiveR; Figure S3: Common qualitative assessments for data quality; Figure S4: Loadings plot from PCA; Figure S5: An example of normalization strategies with label-free quantitative glycoproteomics data from IPX0011732000; Figure S6: Coefficients of variation; Figure S7: Glycoprotein Rank plots; Figure S8: Inspecting overlap in glycoproteomics datasets

## ACKNOWLEDGEMENTS

N.M.R. acknowledges funding from the National Institutes of Health (R00GM147304 and P30CA015704), the Kinship Foundation (Searle Scholar Fellowship), and from the American Association of Cancer Research (Miriam Counts Innovation and Discovery Grant 25-80-39-RILE). GlycoDiveR’s visualizations are inspired by years of work from many people; therefore, we’d like to thank all who contributed to these efforts. We thank the Riley Research group and Shelley Jager for testing GlycoDiveR during the early stages of development.

## FOR TOC ONLY

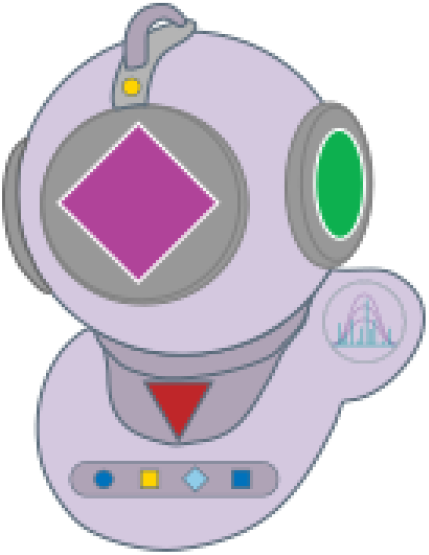

