## Supplemental Information for "GlycoDiveR: a modular R framework to analyze and visualize highly dimensional glycoproteomics data"

**Supplemental Materials**

Figures in this manuscript used data from Kawahara et al. (PXD051882) that were reprocessed with MSFragger-Glyco. Search results that were imported into GlycoDiveR are available as supplemental files:

TMT1_psm.tsv
TMT2_psm.tsv
TMT3_psm.tsv
TMT4_psm.tsv

**Supplemental Figures**

**Figure S1**: Architectural overview of GlycoDiveR
**Figure S2**: Comparing identifications with GlycoDiveR
**Figure S3**: Common qualitative assessments for data quality
**Figure S4**: Loadings plot from PCA
**Figure S5**: An example of normalization strategies with label-free quantitative glycoproteomics data from IPX0011732000
**Figure S6**: Coefficients of variation
**Figure S7**: Glycoprotein Rank plots
**Figure S8**: Inspecting overlap in glycoproteomics datasets

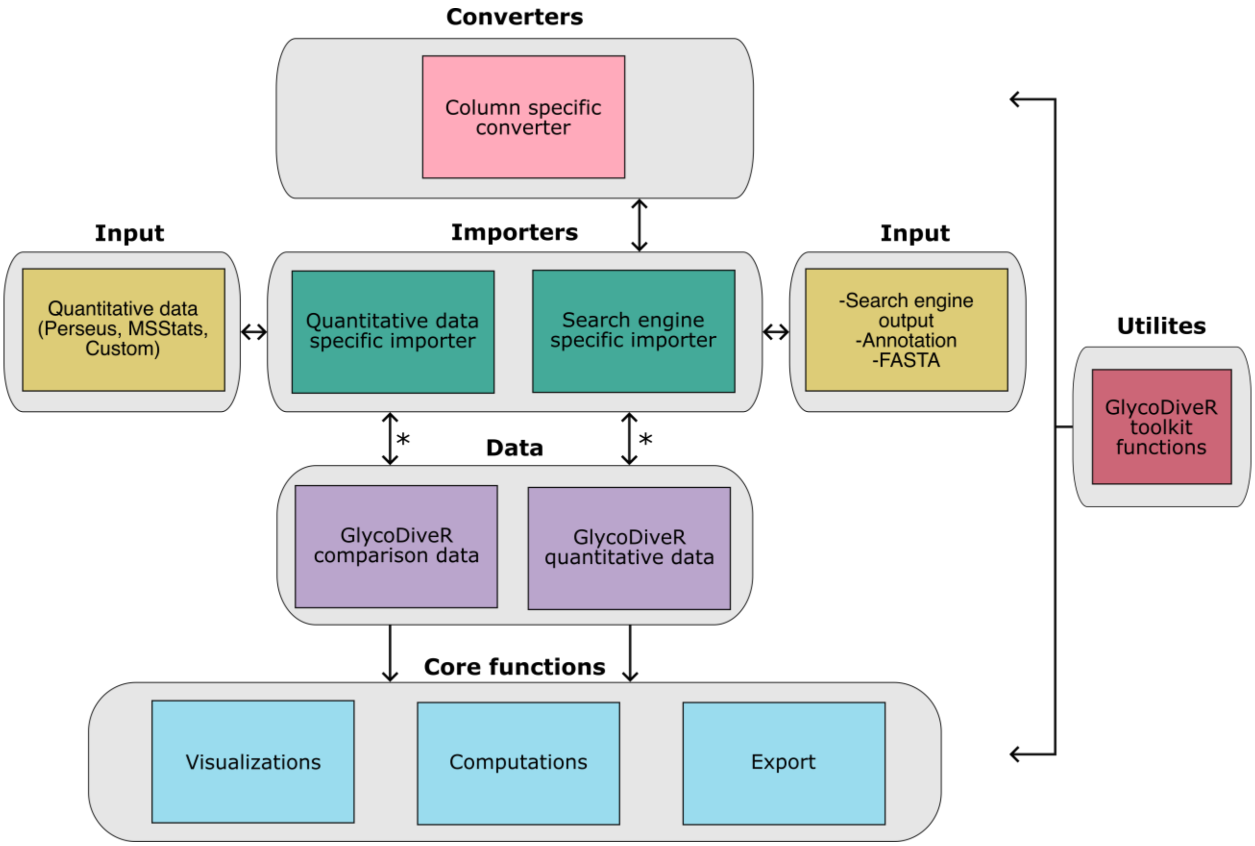


**Figure S1. Architectural overview of GlycoDiveR.** The GlycoDiveR data formats (in purple) are used for all GlycoDiveR visualizations, computations, and exports. A single function call (denoted by *) is used to import and format the data. Each importer is search-engine-specific and has a column-specific converter. Generic functions used throughout GlycoDiveR are stored in the GlycoDiveR toolkit function for clear organization of the framework.

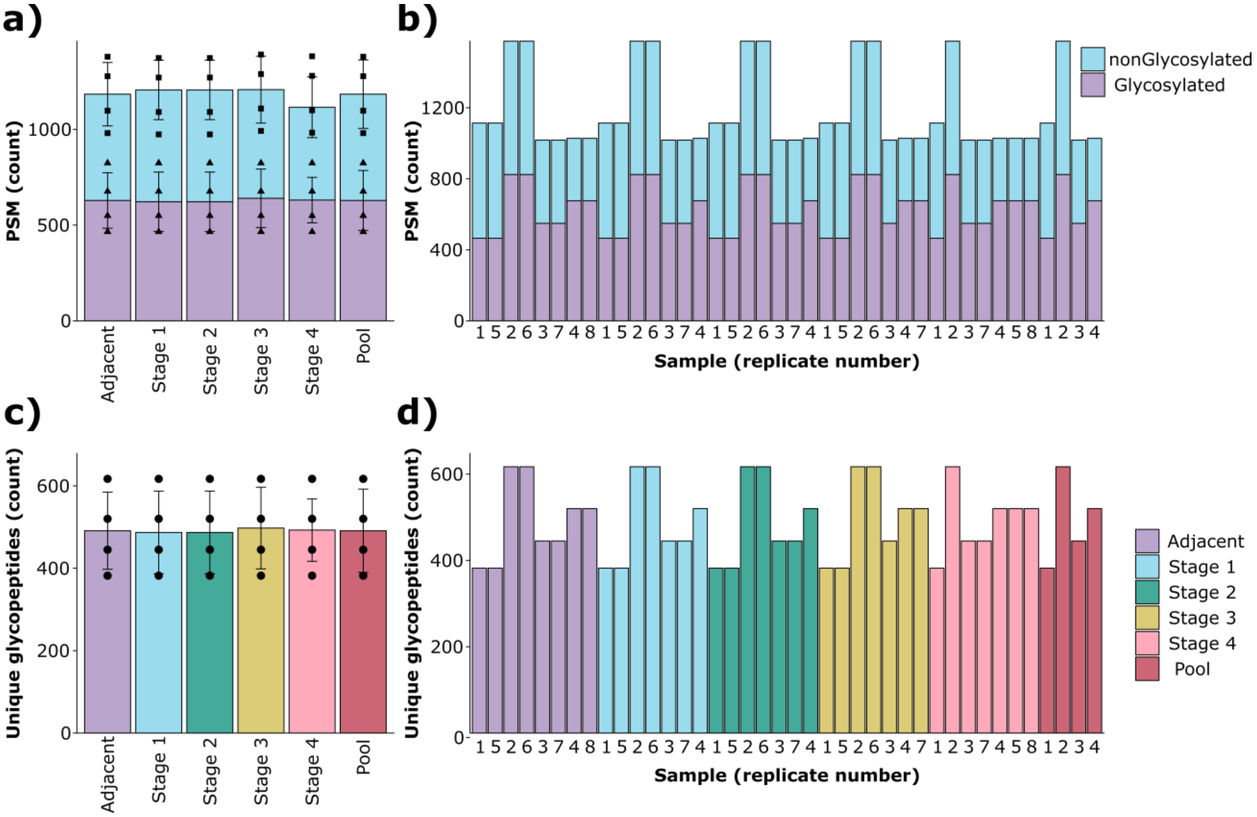


**Figure S2**. **Comparing identifications with GlycoDiveR.** The number of (**a**) total PSMs and (**c**) unique glycopeptides are visualized using either the PlotPSM or PlotGlycopeptide functions . In panels a and c, identifications are grouped per condition, with replicates shown as points on the bar graph and one standard deviation shown as error bars. The same data can visualized with each replicate shown as an individual bar (**b**/**d**) by simply passing the argument *grouping* = *"technicalReps"* to the function.

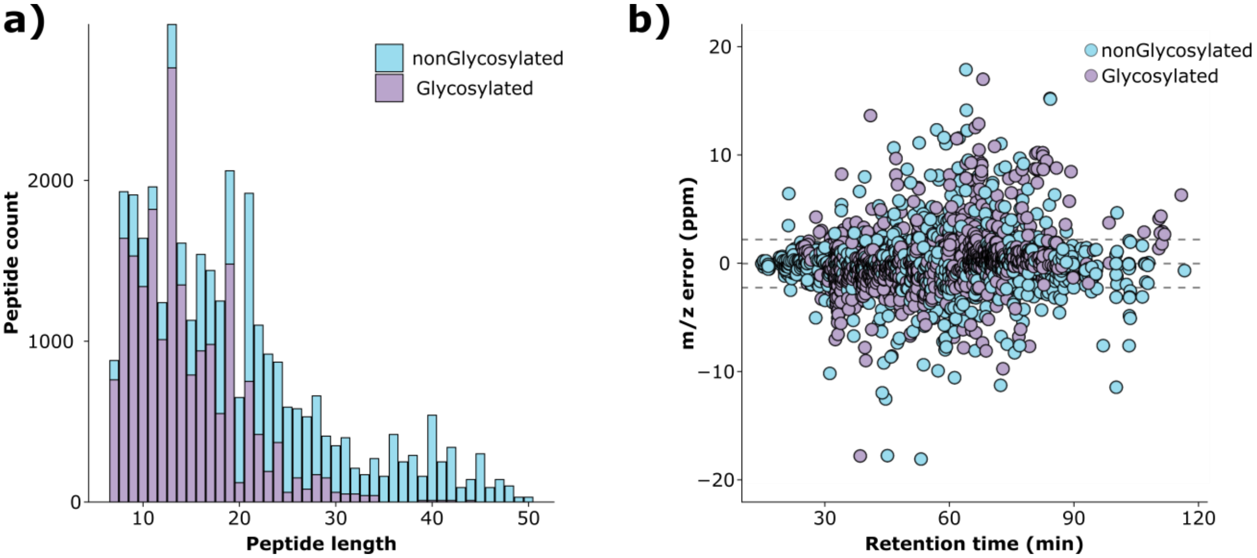


**Figure S3. Common qualitative assessments for data quality.** GlycoDiveR (**a**) generates peptide length distributions for both non-glycosylated and glycopeptides and (**b**) plots mass error (in ppm) versus retention time.

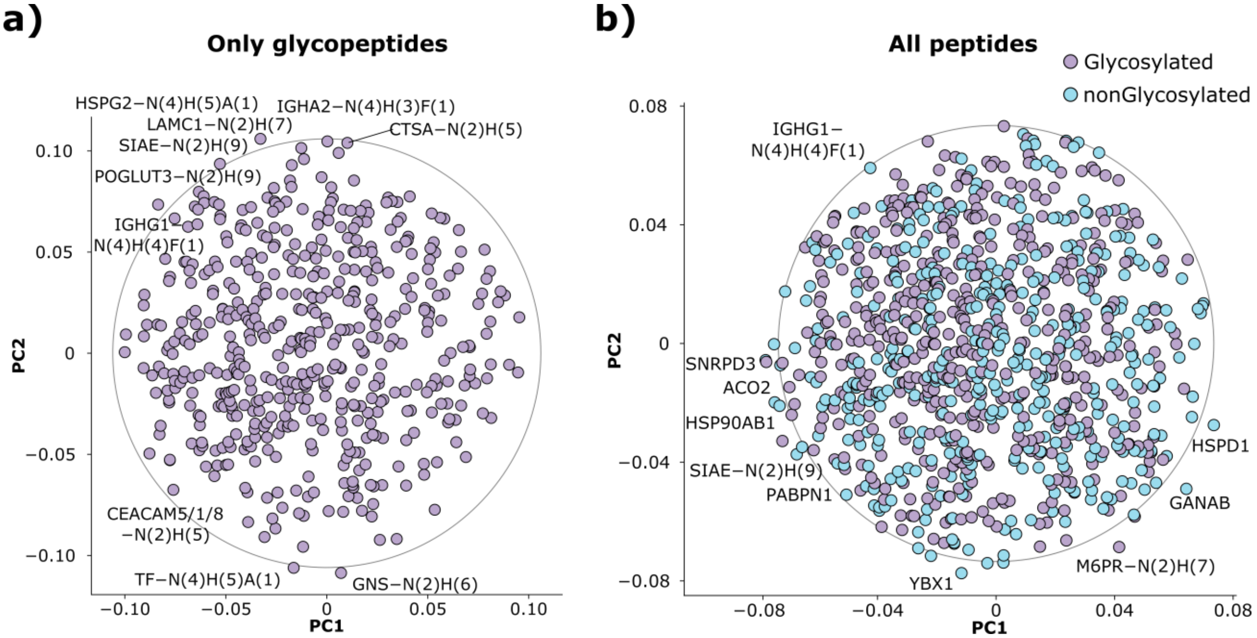


**Figure S4. Loadings plot from PCA.** Examining the major contributors to principal components helps assess which features drive clustering in PCA. Loadings plots are designed to identify these contributors. GlycoDiveR generates loadings plots for PCA plots generated using the PlotPCA function. (**a**) The loadings plot for the Kawahara data, when only glycopeptides are included. (**b**) The loadings plot that also includes non-modified peptides. Both plots provide labels for the ten species that contribute most to separation in the PCA plot.

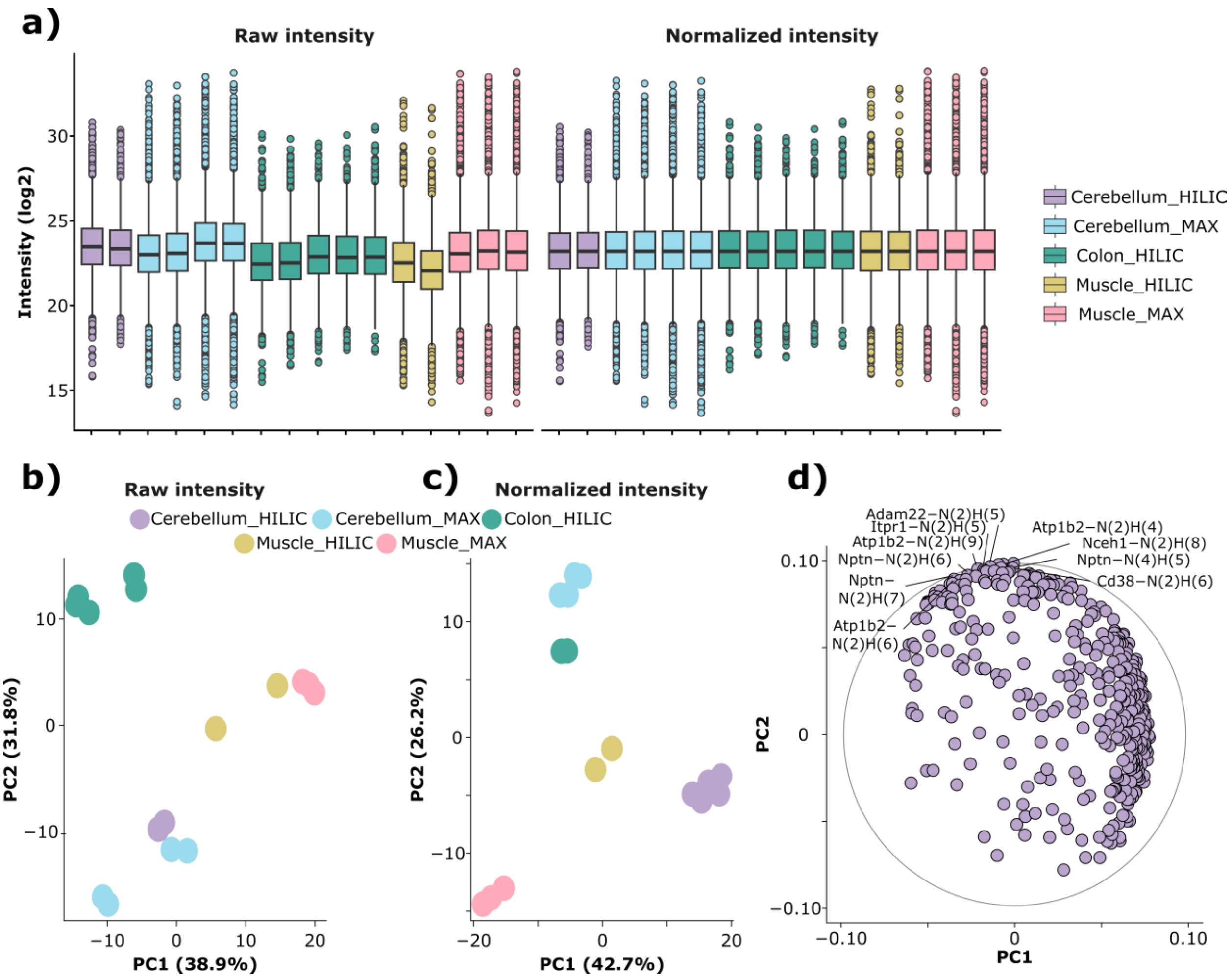


**Figure S5. An example of normalization strategies with label free quantitative glycoproteomics data from IPX0011732000**. (**a**) The PlotAbundanceQC function shows intensity values and the effects of normalization from label free data (instead of TMT data as shown in the main text. The left and right panels show raw intensity and median normalized intensity distributions, respectively. PCA plots for (**b**) raw and (**c**) median normalized intensities and (**d**) a loadings plot for the median normalized intensities are also provided. This represents a common example in quantitative glycoproteomics where median normalization is a reasonable choice for further quantitative processing.

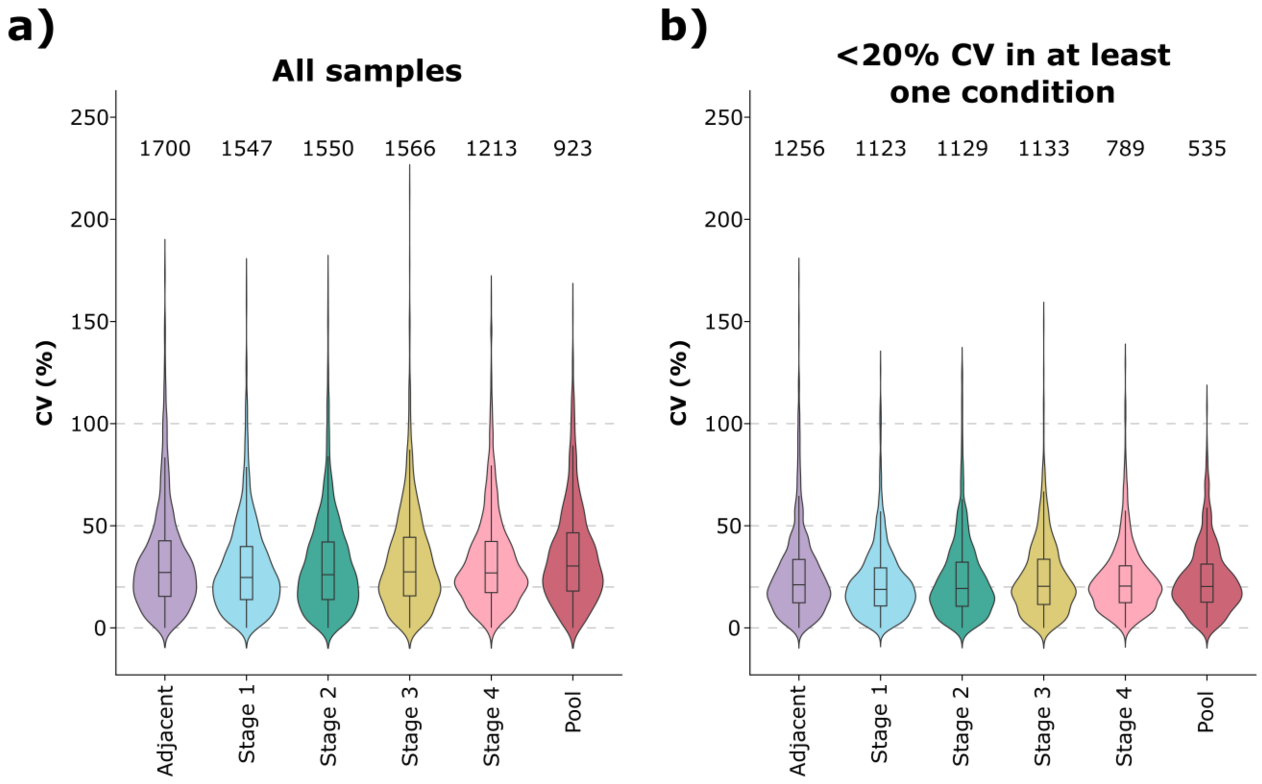


**Figure S6. Coefficients of variation**. GlycoDiveR generates provides sample-specific or whole dataset distributions for glycopeptide coefficients of variation (CVs). Users can choose to retain glycopeptides with CVs below a specified threshold in at least one condition. From example, panels (**a**) and (**b**) show CV distributions for the Kawahara dataset for all glycopeptides, or filtered to retain glycopeptides that were required to have < 20% CV in at least one condition, respectively.

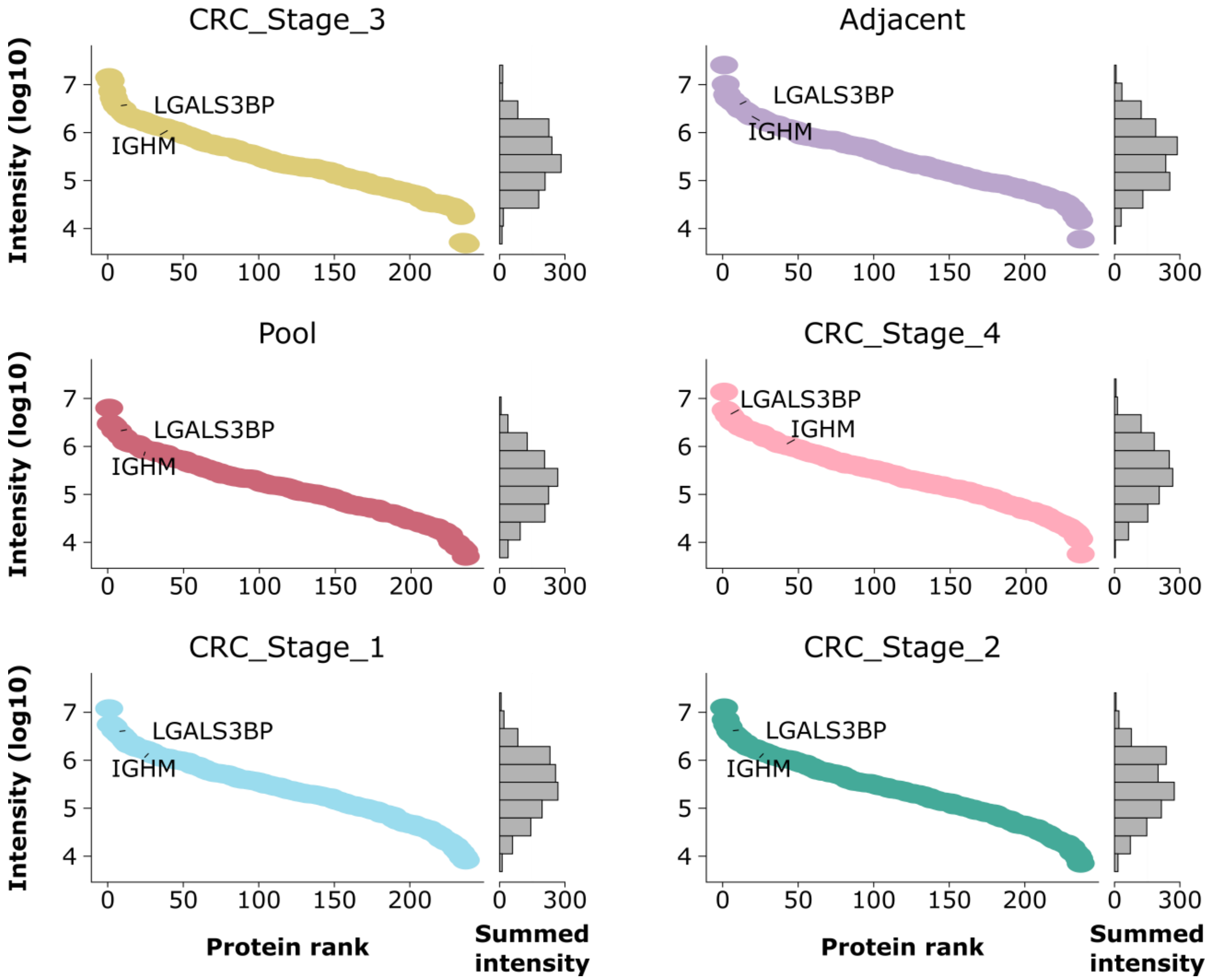


**Figure S7. Glycoprotein Rank plots**. These graphs made by GlycoDiveR plot abundances of glycoproteins (represented by the sum of their glycopeptide intensities) in rank order. The y-axis on the left provides the log10 of summed intensity and the x-axis shows the rank order. The bar graph on the right of each graph shows the distribution of proteins across the intensity scale, i.e., it represents the density of points along the rank curve.

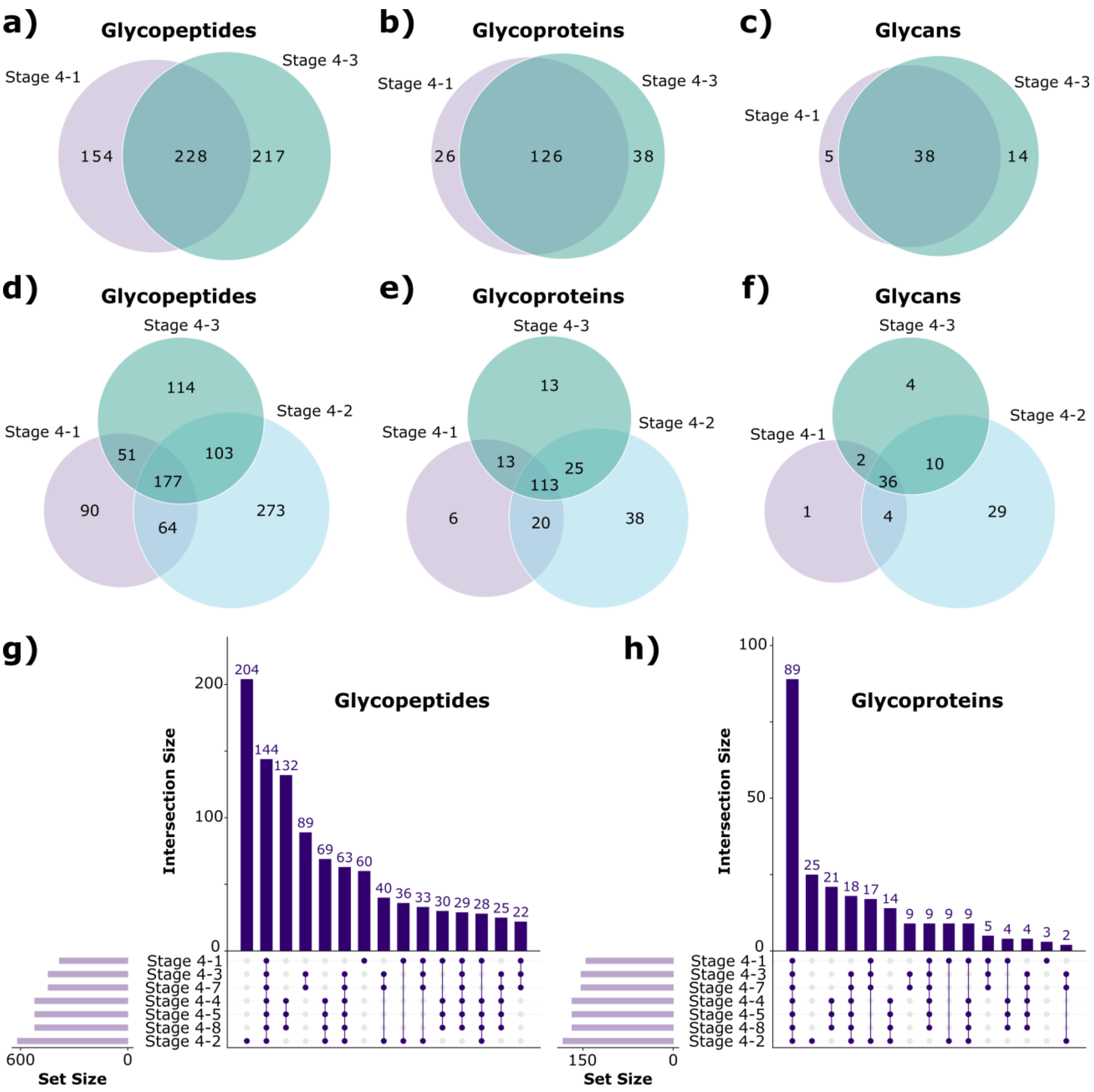


**Figure S8. Inspecting overlap in glycoproteomics datasets.** GlycoDiveR enables Venn diagram comparisons of glycopeptides, glycans, and glycoproteins for up to three conditions. Two-set Venn diagrams are always proportional to the values (**a-c**). Three-set Venn diagrams can be useful for quick visualization (**d-f**),, but they are often mathematically impossible to scale proportionally. To compare three or more sets, GlycoDiveR generates UpSet Plot comparisons (**g-h**), as these can be reliable for quantitatively comparing intersecting data points for any number of conditions.
